# Resurrected protein interaction networks reveal the innovation potential of ancient whole genome duplication

**DOI:** 10.1101/074989

**Authors:** Zhicheng Zhang, Heleen Coenen, Philip Ruelens, Rashmi R. Hazarika, Tareq Al Hindi, Georgianna K. Oguis, Vera van Noort, Koen Geuten

## Abstract

The evolution of plants is characterized by several rounds of ancient whole genome duplication, sometimes closely associated with the origin of large groups of species. A good example is the γ triplication at the origin of core eudicots. Core eudicots comprise about 75% of flowering plants and are characterized by the canalization of reproductive development. To better understand the impact of this genomic event, we studied the protein interaction network of MADS-domain transcription factors, which are key regulators of reproductive development. We accurately inferred, resurrected and tested the interactions of ancestral proteins before and after the γ triplication and directly compared these ancestral networks to the networks of *Arabidopsis* and tomato. We find that the γ triplication generated a dramatically innovated network that strongly rewired through the addition of many new interactions. Many of these interactions were established between paralogous proteins and a new interaction partner, establishing new redundancy. Simulations show that both node and edge addition through the γ triplication were important to maintain modularity in the network. In addition to generating insights into the impact of whole genome duplication and elementary processes involved in network evolution, our data provide a resource for comparative developmental biology in flowering plants.

## Introduction

Comparative analysis of genome sequences, transcriptomes, and phylogenetic and synteny analyses of individual gene lineages placed an ancient hexaploidization event named the gamma (γ) triplication in the stem lineage of core eudicots, before the divergence of Gunnerales and after the divergence of Bu; Jiao et al., 2012; Bowers et al., 2003; Tang et al., 2008). One absolute time estimate for the γ triplication is that it occurred 120 Mya (Figure 1) (Vekemans et al., 2012). The origin of core eudicots marks an important event in plant evolution as today this lineage comprises approximately 75% of all species of flowering plants (Willis and McElwain, 2013; Soltis et al., 2003). Aside from the γ triplication and the presence of ellagic and gallic acid, the group shares few unique characteristics (Stevens and Davis, 2006). However, the Pentapetalae, which comprise most core eudicots but originated a few million years later, are morphologically more distinct and are characterized by the ‘canalization’ or a more clear definition of flower development (Theißen et al., 2016a; Waddington, 1942; Melzer et al., 2016; Soltis et al., 2003). In this group, floral organs are in pentamerous whorls and a clear separation of sepal and petal identity exists (Soltis et al., 2003; Stevens and Davis, 2006). Therefore, while core eudicots share the γ triplication, it appears that the morphological consequences of this genomic event were established only somewhat later in evolution and are more apparent from Pentapetalae onwards (Vekemans et al., 2012; Schranz et al., 2012). In the context of developmental genetics, the origin of Pentapetalae has been proposed to coincide with a transition from a fading borders model with overlapping gene expression domains of floral organ identity genes, to an ABCDE model with more strictly defined expression domains (Chanderbali et al., 2016; Soltis et al., 2009; Soltis and Soltis, 2016).

**Figure 1.**
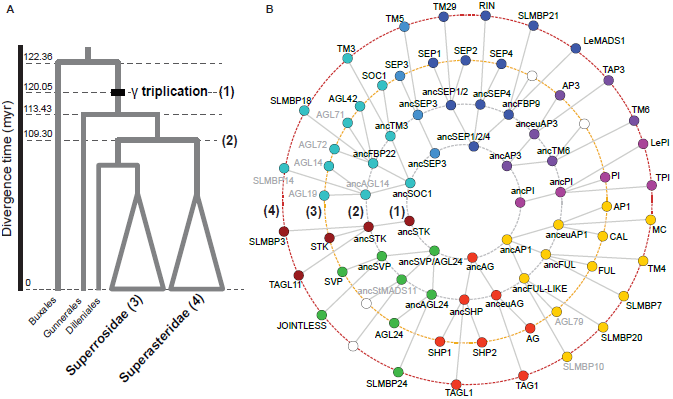
Evolution of selected MADS-box proteins of interest. (A) Simplified phylogenetic tree of eudicots. The positions at which protein interaction networks were inferred are indicated; (1) PIN just before γ triplication or Pre-PIN, (2) PIN at the Asterid-Rosid split or Post-PIN, (3) PIN of *Arabidopsis thaliana* (Ara-PIN) and (4) PIN of *Solanum lycopersicum* (Sol-PIN). (B) Phylogenetic relationships between ancestral and extant proteins present in each reconstructed PIN. Reconstructed ancestral proteins are named after their *Arabidopsis* or tomato descendants preceded by ‘anc’ for ancestral. Gray font proteins were not included in this study (see methods). White circles indicate the absence of a specific protein due to gene loss. Abbreviations: AP: APETALLA. FUL: FRUITFULL. FUL-LIKE: FRUITFULL-like gene. CAL: CAULIFLOWER. MC: MACROCALYX. SLMBP: *Solanum lycopersicum* MADS-box protein. DEF: DEFICIENS. TM: Tomato MADS-box gene. TAP: Tomato APETALLA. GLO: GLOBOSA. PI: PISTILLATA. AG: AGAMOUS. SHP: SHATTERPROOF. TAG: Tomato AGAMOUS. TAGL: Tomato AGAMOUS-like gene. STK: SEEDSTICK. SEP: SEPALLATA. FBP: FLORAL-BINDING PROTEIN. SVP: SHORT VEGETATIVE PHASE. SOC1: SUPPRESSOR OF OVEREXPRESSION OF CONSTANS 1

The duplication patterns of MADS-domain proteins 舒 a conserved class of transcription factors that act as key regulators of reproductive development in flowering plants 舒 indicate that many gene lineages present in extant core eudicots are derived from this whole genome triplication with most being retained in duplicate or triplicate copies (Vekemans et al., 2012; Viaene et al., 2010; Kramer, 2004; Hernásuccess of core eudicotsndez-Hernández et al., 2007; Airoldi and Brendan, 2012; Litt and Irish, 2003; Kramer et al., 2006). Their molecular function as transcription factors requires them to localize in the nucleus and form specific multimeric transcriptional complexes to regulate downstream targets (Smaczniak et al., 2012; Immink et al., 2010). Considering the critical role of the specific protein binding affinities among these proteins in the induction of flowering, inflorescence meristem specification, floral meristem and floral organ specification, the expansion of MADS-box genes through the γ triplication and the protein interactions that evolved may have played an important role in establishing the derived morphology of Pentapetalae and the success of core eudicots (Theißen et al., 2016b; Liu et al., 2010; Veron et al., 2007; Shan et al., 2009). The functional importance of protein interactions of MADS-domain proteins has been characterized through genetic analysis in *Arabidopsis thaliana* (Liu et al., 2009a; Smaczniak et al., 2012; Yan et al., 2016) and comprehensive yeast two-hybrid protein interaction maps for MADS-domain proteins are available for this model system and a few other species (Leseberg et al., 2008; de Folter, 2005a; Immink et al., 2009; Angenent and Immink, 2009; Ruokolainen et al., 2010). While such data allow to trace the origin and evolutionary diversification of protein interactions and some of their functions, such inferences suffer from sparse sampling and different yeast assays being used, which hampers direct comparison of data and consequently the accuracy of deep evolutionary inferences.

Biological networks are characterized by several organizational properties to which certain biological advantages can be attributed. The most often used property of nodes in a network is the degree, or the number of interactions of a protein in a protein interaction network. The degree distribution of networks is usually heterogeneous or mathematically scale free, with few nodes having many interactions and many nodes having few (Barabasi and Albert, 1999a). This property indicates the presence of hubs in the network, or very well-connected nodes. The origin of this property of the network is closely linked to its origin through gene duplication as more connected nodes will acquire more interactions through duplication, a mechanism referred to as preferential attachment (Eisenberg et al., 2003). The presence of hubs in a network is considered to make the network more robust to random failure, as the small number of hubs decreases the likelihood of these being affected. Another important property is the degree of clustering or modularity. Modularity and also hierarchy ߞ modularity of modules ߞ are considered to originate from a cost associated with connections between nodes (Clune et al., 2013; Mengistu et al., 2016). The evolutionary advantage of a modular organization is that it makes the network more adaptable as modules can be easily added or removed (Tran and Kwon, 2013; Bassett et al., 2010; Mengistu et al., 2016).

Specifically in plants, but of general biological importance, the role of massively concerted gene duplications at the genome level is well documented (Debodt et al., 2005; Adams and Wendel, 2005; van Hoek and Hogeweg, 2009; Conant and Wolfe, 2006; Arabidopsis Interactome Mapping Consortium et al., 2011; Soltis and Soltis, 2016). Such whole-genome duplication events could also have a major effect on the rewiring of protein interaction networks as predicted by the duplication-divergence model (Wagner, 2001; De Smet and Van de Peer, 2012; Arabidopsis Interactome Mapping Consortium et al., 2011). To understand the impact of the γ triplication on the origin of a large group of plant species, we studied the intricate ancestral protein interaction networks of MIKC^C^ MADS-domain transcription factors. We resurrected ancestral proteins immediately before this genome triplication and 10 million years later, at the diversification of rosid and asterid flowering plants. By directly comparing ancestral networks with extant networks from *Arabidopsis* and tomato in a single experimental setup, we are able to go beyond theoretical models and comparative analysis of present-day networks and instead pinpoint directly how this network diverged and which processes were responsible for its origin and divergence.

## Results

### Accurately resurrected ancestral MADS-domain proteins reveal the origin of extant networks

We reconstructed protein interaction networks (PINs) between representatives of nine MADS-box gene subfamilies at three distinct points in time for a total of four PINs: (1) just before the γ triplication event coinciding with the origin of the core-eudicots, (2) following the γ triplication at the Asterid-Rosid split and present-day (3) from *Arabidopsis thaliana* and (4) *Solanum lycopersicum* (tomato) (Figure 1). These ancestral and extant interaction networks are respectively referred to as Pre-PIN, Post-PIN, Ara-PIN and Sol-PIN throughout this study.

Reconstruction of the ancestral proteins that form the Pre-PIN and Post-PIN was performed by inferring the maximum-likelihood protein sequences at the ancestral nodes of interest for each subfamily separately (see methods, Supplemental Figure 1). MADS-box genes are a good model system for ancestral sequence resurrection since there are many sequences available throughout the angiosperm phylogeny which are mostly well conserved within their subfamilies. The accuracy of reconstructed ancestral protein sequences is represented by the posterior probability of each inferred amino acid in the ancestral sequence (Supplemental Figure 2). Both the ancestral proteins before and after γ triplication were reconstructed with on average 92.8% and 94.6% of sites obtaining a posterior probability higher than 0.95. Previous studies utilizing ancestral proteins to characterize evolutionary transitions defined ambiguously reconstructed sites as those sites for which the most likely amino acid has a posterior probability lower than 0.80 and that have an alternative amino acid state with a posterior probability higher than 0.2 (Voordeckers et al., 2012; McKeown et al., 2014; Anderson et al., 2016; Eick et al., 2012). Surveying ambiguous sites in the ancestral proteins reconstructed in this study revealed that ambiguously inferred sites were mostly located outside of the highly conserved K-domain (Supplemental Figure 2), which plays a prominent role in interactions between MADS-box proteins (Yang and Jack, 2004; Fan et al., 1997; Kaufmann et al., 2005). Out of the 26 reconstructed ancestral proteins, only 11 ambiguous sites in the K-domain had an alternative amino acid not biochemically similar to the most likely amino acid. Given the scale of our study, this represents only 0.5% of all reconstructed sites in the K-domain. Following inference, codon optimized ancestral DNA sequences were synthesized and, analogous to their *Arabidopsis* and *Solanum* descendants, cloned into yeast two-hybrid (Y2H) expression vectors. Subsequently, all pairwise interactions for each set of MADS-box protein constructs at an ancestral or extant node were determined using a high-throughput yeast two-hybrid system (Figure 2). In total 582 binary protein-protein interactions were tested (Supplemental Dataset 3).

**Figure 2.**
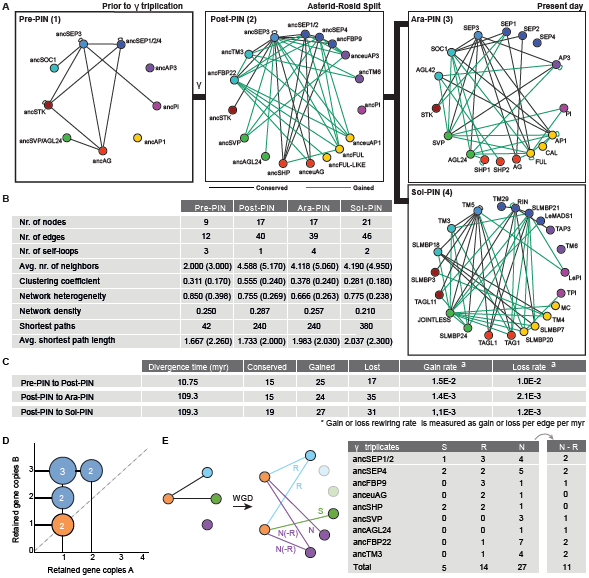
Evolution of MADS-box protein interaction networks reveals accelerated rewiring following γ triplication. (A) MADS-box protein interaction networks just before the γ triplication (1), at the Asterid-Rosid split (2), for Arabidopsis (3) and for Solanum (4) as assayed by Y2H. Same color nodes indicate homologous MADS-box gene lineages as mentioned in Figure 1B. Conserved and gained protein interactions are indicated as black and green solid lines respectively, and are determined by comparing Post-PIN to Pre-PIN and Ara-PIN or Sol-PIN to Post-PIN. (B) Network topological measurements for reconstructed Y2H MADS-box PINs. Values between parentheses denote the average network topological measurements of 10,000 randomly generated networks of same network size and average degree. (C) The rate of interaction gain and loss as determined from PrePIN to Post-PIN and from Post-PIN to Ara-PIN or Sol-PIN for Y2H. Gain and loss rate was defined as gained or lost edges divided by number of potential interactions (selfloops included) times the divergence time. (D) According to the gene dosage balance hypothesis, the proteins (A and B) involved in the nine PPIs observed in Pre-PIN Y2H should be retained in similar balance after the γ triplication. The graph shows the dosage of the gene copies of these proteins. The number of interacting proteins that were retained in balance after ? are marked in orange. (E) Schematic overview of the duplication-divergence possibilities as observed in the Y2H PINs and number of interactions that show: R, Redundancy; S, subfunctionalization or N, neofunctionalization when we compare their interactions from the Pre-PIN to the Post-PIN. N-R or neo-redundant interactions are those neofunctionalized interactions that redundantly interact with the same paralogs. Genes that did not duplicate or had no interactions in Pre-PIN are excluded.

While genetic evidence has supported the functional importance of the protein interactions of MADS-domain as determined by yeast-two-hybrid (Y2H), the method is prone to false positives and dependent on the yeast strain and vector system being used (Chen et al., 2010). Therefore, we first determined the reliability of the Y2H system used in this study and consequently our interaction data. In the absence of a curated interaction data set for MADS-box proteins, we compared Ara-PIN and Sol-PIN to *Arabidopsis* and *Solanum* MADS-box protein interaction networks described in the literature (Supplemental Figure 3) (de Folter, 2005a; Leseberg et al., 2008). Our analysis shows a high overlap between the Ara-PIN and Sol-PIN described here and previously constructed networks. We determined the accuracy or similarity of Ara-PIN and Sol-PIN to be 0.80 and 0.74 respectively. The overall similarity between both networks is 0.77 (Supplemental Figure 3). The overall high similarity between Ara-PIN or Sol-PIN and previously described *Arabidopsis* and *Solanum* MADS-box protein interaction networks strengthens our confidence in the ancestral Pre-PIN and Post-PIN networks.

**Figure 3.**
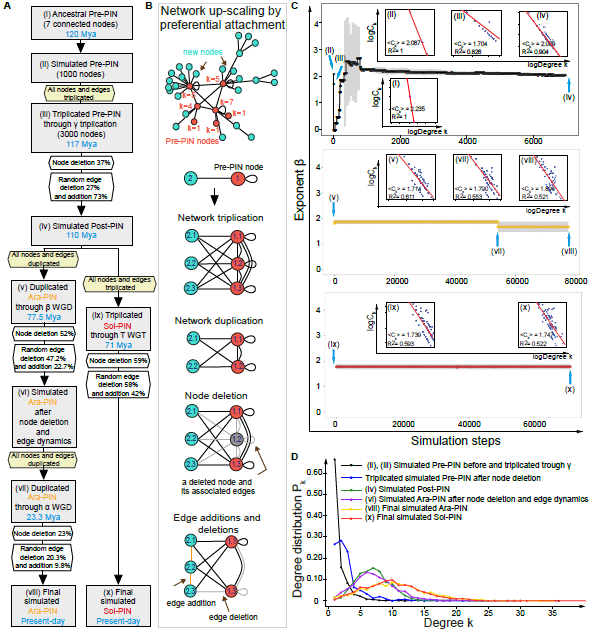
Tracing elementary processes in network evolution from Pre-PIN until extant Ara-PIN and Sol-PIN. (A) The flowchart shows the initial up-scaled Pre-PIN model with the simulation process based on the actual ancestral and extant MADS-box PIN parameters (calculated percentages of node deletion, edge addition and deletion). The initializing network is up-scaled to the size of 1000 nodes based on the actual Pre-PIN with 12 edges and 7 connected nodes where 2 isolated nodes are excluded. Each simulation step reflects bins of 0.001 myr for random edge addition or deletion excluding the steps of ancient whole genome triplication or duplication and node deletions. (B) Elementary processes of network evolution. The mechanisms of network up-scaling using preferential attachment in the initial upscaled Pre-PIN model, network triplication, node deletion and edge additions and deletions applied in the simulated networks. The ancestral Pre-PIN nodes are labelled in red while the new nodes are labelled in turquoise. All nodes are numbered and the number near the Pre-PIN nodes indicate their corresponding node degrees. (C) Exponent (3 distribution for the initial up-scaled Pre-PIN model. The main plots show the average value of the (3 exponent together with the standard deviation (plots in black, yellow and red and error bars in grey) for a total 10 replicated simulations. Blue arrows indicate positions of several important simulated networks following the simulation steps corresponding to the simulation process from A. There are total of nine intrinsic log-log plots showing the relationship between degree *k* and *C_k_* of each node in simulated MADS-box PINs following the simulation steps (for the complete plots, see Supplemental Figure 4). (D) Degree *P_k_* distribution for the initial up-scaled Pre-PIN model.

### Node expansion after □ of MADS-domain proteins is not dosage balanced

Throughout plant evolution, series of whole-genome duplications have expanded the number of nodes in major regulatory networks including the MADS-box gene family (Veron et al., 2007; Vekemans et al., 2012; Ruelens et al., 2013; Jiao et al., 2012, 2011). Following the γ hexaploidization event, theoretically all genes were triplicated, however, quickly afterwards redundant genes would be silenced and lost through pseudogenization (Wendel et al., 2016; Freeling et al., 2012). Indeed, from Pre-PIN to Post-PIN not all genes were retained in three copies, generating an ancestral Post-PIN with a network size only two fold larger than the original Pre-PIN (Figure 2A) (Veron et al., 2007). For proteins that function in multimeric complexes, gene retention after whole genome duplication is often explained by the dosage balance hypothesis, which states that specific types of interacting proteins, such as transcription factors, are retained in similar dosage not to disturb dosage-sensitive processes (Veitia and Potier, 2015; Edger and Pires, 2009). Since our approach provides us with ancestral interactions, it is possible that we would observe this process directly. To investigate dosage balance, we plotted post γ gene dosages of the proteins that interacted before γ by Y2H (Figure 2D). However, in line with previous findings (Guo et al., 2013), we did not observe dosage-balanced gene retention in MADS-networks as only two out of nine proteins are retained in balance. After the Asterid-Rosid split, two additional rounds of ancient whole genome duplications occurred along the lineage towards Arabidopsis (*β*, 77.5 Mya and *α*, 23.3 Mya) and one ancient whole genome triplication towards *Solanum* (T, 71 Mya) (Bowers et al., 2003; Tomato Genome Consortium, 2012). Here, the expansion of the network was limited as only 22 homologous proteins are present in *Arabidopsis* and 23 in *Solanum* as compared to 19 proteins post γ triplication (Figure 1B), illustrating that unlike the γ triplication, the more recent ancient whole genome duplications did not result in the strong expansion of the MADS-box PIN.

### Strong rewiring, neofunctionalization and neoredundancy after hexaploidization

The γ triplication innovated the MADS-domain protein network by the addition of new nodes, yet duplication is not the only force driving changes of PINs. Edges or interactions can be gained, lost and rewired and as a consequence the functional information in the network could have evolved (Pastor-Satorras et al., 2003; Vázquez et al., 2002; De Smet and Van de Peer, 2012).

We noticed that the interaction patterns of the post-PIN were much more similar to those of Ara-PIN and Sol-PIN as compared to Pre-PIN, even though they are divided by 110 mya of evolution as compared to 10 mya between Pre-PIN and Post-PIN. To investigate whether edge rewiring happened at a faster rate following genome duplication, we calculated and compared the average rate of interaction gain and loss from Pre-PIN to Post-PIN with the rates from Post-PIN to Ara-PIN and Post-PIN to Sol-PIN (Figure 2C). Our results show that from Pre-to Post-PIN the rate in interaction gain is 1.5E-02 gained edges per total possible edges per myr, approximately 1.5 fold higher than the rate of interaction loss (1.0E-02/edge/myr). In addition, from Post-PIN to Ara-and Sol-PIN new interactions evolved at a rate of 1.4E-03 and 1.1E-03/edge/myr respectively, while interactions were lost at 2.1E-03 and 1.2E-03/edge/myr, respectively. These results indicate that in the 10 myr following the γ triplication, the MADS-box network rewired at a rate approximately tenfold higher than over the 110 myr between Post-PIN and Ara-PIN or Sol-PIN.

Moreover, from the origin of core eudicots up to the Asterid-Rosid split, the network rewired mainly through the gain of new interactions. By contrast, from the Asterid-Rosid split until present-day, not only did the overall rewiring rate decrease, interaction loss became higher than interaction gain (Figure 2C). Together, these results indicate that shortly following the γ triplication, the MADS-box PIN underwent accelerated rewiring.

The γ triplication added many new interactions to the network, which may have had consequences for the selectivity with which these interactions could be maintained. To understand this specificity, it is sensible to compare network density, i.e. the ratio between the number of actually observed interactions and the number of possible interactions. Despite strong edge addition, network density did not notably change from Pre-PIN to the current PINs, Ara-PIN and Sol-PIN (Figure 2B). This relatively constant density suggests that an optimal number of specific interactions can be maintained by a set number of MADS-domain proteins, a property that probably relates to protein structure (Zarrinpar et al., 2003).

Rewiring can be a consequence of neofunctionalization or subfunctionalization (He and Zhang, 2005). When applied to protein interactions, the neofunctionalization model implicates that following duplication, one paralog retains all interactions while the other is released from functional constraints and undergoes rapid diversification, resulting in the evolution of novel interactions. The subfunctionalization model posits that paralogous proteins rewire by redistributing their ancestral interactions among the different paralogs without new interactions emerging. In agreement with the rapid rewiring after the γ triplication, our data show many more instances of neofunctionalization than of subfunctionalization when comparing Pre-PIN to Post-PIN or Post-PIN to Ara-PIN and Sol-PIN. Rewiring following the γ triplication, however, can not be explained by a strict interpretation of sub-or neofunctionalization (Figure 2E) (Voordeckers et al., 2012). Rather, the data show both rapid rewiring of all descendant paralogs by acquiring novel interactions and a combination of sub-and neofunctionalization, in which paralogs both acquire new interactions while maintaining a set of ancestral interactions.

Interestingly, we observe many cases in which new redundancy originated through the γ triplication; i.e two newly emerged paralogs interact with the same protein while their ancestor did not (Figure 2E). This observation, which we refer to as neoredundancy, can be explained by the fact that new paralogs are highly similar and as a consequence a protein evolving to interact with one of these paralogs will likely also interact with the other paralog. Together, our data suggest that the γ triplication dramatically innovated the MADS-PIN, but at the same time the network also acquired novel robustness through the redundancy that was established.

### Ancestral and extant networks are heterogeneous, small world and modular

We observed that the γ triplication duplicated the number of nodes and rewired interactions. How these edges are mathematically organized in the network is referred to as the topology of the network. The presence of hubs, the number of modules and the organization of modules all potentially contribute to network robustness and evolvability (Lachowiec et al., 2016; Clune et al., 2013; Mengistu et al., 2016). To understand the effect of ancient hexaploidization on the topology of the network, we calculated a number of topological network parameters commonly used to describe networks. For comparison, we also determined these parameters for random networks of equivalent size and average degree (Figure 2B).

A highly heterogeneous degree distribution is suggestive of the presence of hubs in a network as hubs have many connections while other proteins have only a few connections. By contrast, in random networks all nodes have approximately the same number of connections, with only a small deviation (Albert et al., 2002). As compared to random networks, all MADS-box PINs exhibit a high network heterogeneity (Figure 2B). Indeed, in Pre-and Post-PIN more than 40% of all connections involve SEP3 proteins with a degree of 7 and 16, approximately threefold the network average degree of 2.6 and 4.7 respectively. However, both in Ara-PIN and Sol-PIN, there is no single prominent node with a high maximum degree. Rather, in the latter two networks, multiple proteins exhibit a moderately high degree between 8 and 12, only two-fold the average degree. These new hubs are FUL, AP1, SEP3, SOC1 and SVP in Ara-PIN and JOINTLESS, TM5, RIN and SLMPB21 in Sol-PIN. SEP3 therefore lost its prominence as the sole hub protein because multiple hubs evolved in the lineages towards Ara-PIN and Sol-PIN.

To further understand how the γ triplication affected the topology of the network, we investigated the evolution of clustering or modularity in the network. The extent of clustering is described by the average clustering coefficient of a network, which for most real-world networks is higher than comparable random networks and indicates their modular structure (Albert et al., 2002). For the MADS-box PINs, all Y2H networks have a higher average clustering coefficient than their corresponding random networks (Figure 2B). Moreover, the average clustering coefficient increases following the γ triplication from C_*Pre-PIN*_ = 0.311 to C_*Post-PIN*_ = 0.555, while both descendant networks, Ara-PIN and Sol-PIN, again have a lower clustering coefficient of C_*Ara-PIN*_ = 0.378 and C_*Sol-PIN*_ = 0.281. A clear negative correlation between clustering coefficient and degree can be observed for post-PIN, which is also present in pre-PIN, but such a clear correlation is lost in the extant networks of *Arabidopsis* and tomato (Supplemental Figure 4C). This indicates hierarchy in the ancestral networks, or a modular organization of modules. This modular and hierarchical organisation is considered to originate from a cost associated with individual interactions, which is consistent with the relatively constant density of the networks and is thought to confer evolvability to the networks (Clune et al., 2013; Mengistu et al., 2016).

**Figure 4.**
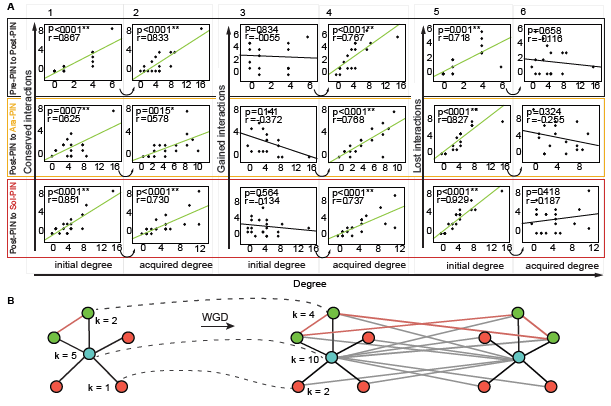
Analysis of MADS-box PINs edge dynamics. (A) Scatter plots indicating node degree versus conserved interactions (1 *and* 2), gained interactions (3 *and* 4) and lost interactions (5 *and* 6) as determined between Pre-PIN and Post-PIN (top), between Post-PIN and Ara-PIN *(middle)* and between Post-PIN and Sol-PIN *(bottom)*. Correlations are represented by the Pearson correlation coefficients of the relationships and the associated p value. (B) Example of preferential attachment as a result of WGD. k is the degree of the proteins before (left) and after (right) a WGD.

In addition to a high clustering coefficient, real-world networks also have a short average path length. The average shortest path length is defined as the average minimal number of edges that connect all possible pairs of nodes in a network. A short average path length ensures an efficient and fast transmission of information throughout the network (Watts and Strogatz, 1998; Barabási et al., 2004). Networks that have a higher clustering coefficient than a comparable random network, yet also have an average shortest path length similar to random networks are referred to as small world networks. The average shortest path length of each Y2H MADS-box PIN was consistently smaller, albeit only slightly, than their random networks and remained relatively stable throughout evolution (Figure 2B). Together with their high clustering coefficient, this indicates that all Y2H PINs meet the requirements of small world networks. Overall we find that the γ triplication did not establish a qualitatively different topological organization of the network and that the post-PIN network evolved from being organized around the single hub SEP3, to a network organized around multiple hubs in *Arabidopsis* and tomato.

### The □ triplication maintained hierarchical modularity through edge and node dynamics

Because the network dynamics of the γ triplication did not disrupt network topology, we asked how network topology was maintained despite extensive rewiring. To statistically evaluate the role of elementary processes that were applied to the network, we applied the observed network dynamics between ancestral and extant networks to simulated large scale networks (Figure 3). An initial large-scale Pre-PIN network was obtained by upscaling the ancestral Pre-PIN of 7 connected nodes to 1,000 nodes by preferential attachment (Figure 3B and Figure 3C, (i) to (ii)). Thereafter, the up-scaled Pre-PIN was subjected to network triplication, node deletions, edge deletions and additions as schematized in Figure 3A. To understand the role of individual elementary processes, we implemented modifications of the simulations and for comparison, we tested the effect of the measured network dynamics on a completely random network model obtained via random attachment of nodes (see Methods) (Supplemental Figure 4). For each simulation, we carried out at least 10 stochastic runs.

Our focus was on how elementary processes contributed to hierarchy in the network, which is measured by a significant negative linear correlation between clustering coefficient *C_k_* and degree *k* (plots in Figure 3C). Triplication of the nodes in the upscaled Pre-PIN did not significantly affect network hierarchy (Figure 3C, (iii)). However, we see that the observed 37% node deletion subsequent to node triplication would have destroyed network hierarchy without edge dynamics (Figure 3C, above *β* exponent distribution). Edge dynamics played an important role in restoring hierarchy in the simulated Pre-PIN (Figure 3C, (iii) to (iv)). The approximately threefold edge addition frequency (73%) compared to edge deletion frequency (27%) facilitated the generation of numerous novel clusters in the simulated Post-PIN, while not many such newly created clusters were eliminated (Figure 3C, (ii) to (iv)). This hierarchy seems subsequently to be retained from simulated Post-PIN to Ara-PIN (Figure 3C, (v) to (viii)) and simulated Sol-PIN (Figure 3C, (ix) to (x)) even though the edge deletion frequency was found 2 and 1.4 fold higher than the edge addition frequency respectively (Figure 3A). A high scaling *β* exponent was obtained (Figure 3C, middle and bottom *β* exponent distribution), which is consistent with earlier studies where several biological networks such as yeast PINs have exhibited higher scaling exponents (Koonin et al., 2007).

We further evaluated the role of the node triplication in network evolution by constructing networks devoid of this major event. In this simulation the network size from Pre-PIN to Post-PIN was retained without triplication. Here, node deletion followed by edge dynamics created clusters in the simulated networks but never hierarchy (Supplemental Figures 5C, 5E and 5G). Indeed, the effect of the edge dynamics was too drastic on such a reduced network size. Therefore the combination of node and edge dynamics appears to have been necessary to maintain hierarchy in the network after the γ triplication (Figure 3C, above *β* exponent distribution). This suggests that modularity was actively maintained through edge and node dynamics. While hierarchical modularity provides the biological advantage of modules that can be easily added or removed, it could also be that the cost associated with the interactions drove the network to retain its hierarchy (Mengistu et al., 2016). Furthermore, because the γ triplication already established many clusters in the network, subsequent whole genome duplications observed in the lineages towards *Arabidopsis* (α WGD and β WGD) and Tomato (T WGT) did not significantly affect hierarchy of the network anymore (Figure 3C, (v) and (vii)). While hierarchy was not clear for Ara-PIN and Sol-PIN in the smaller experimental networks, when upscaled through preferential attachment to a 1000 nodes, also these networks display hierarchy (Supplemental Figure 4C).

**Figure 5.**
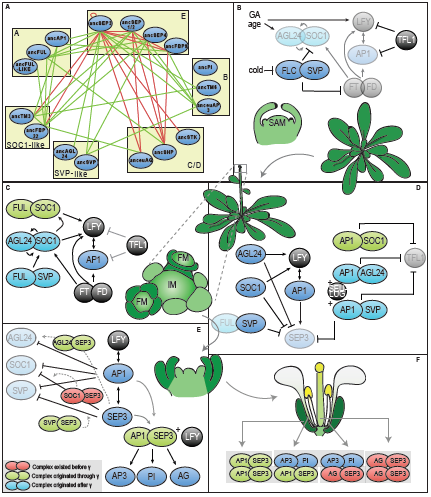
Overview of all MADS-box complexes involved in floral transition in *Arabidopsis*. (A) Post-PIN network. Green lines denote all interactions that originated through, red lines denote interactions that existed before the triplication. Proteins are ordered into their subfamilies by yellow boxes. (B) MADS-domain complexes before (C) and during early floral transition. (D) MADS-domain complexes at the start of floral organ development (stage 1 and 2), (E) during stage 3 and (F) in the mature flower. Red complexes existed before the γ triplication, green complexes evolved through the γ triplication and light blue complexes arose later. Dark blue ellipses are proteins not in complex or not discussed here. Black circles are non-MADS-domain proteins. Transparent proteins are weaker expressed in that stage. Dotted arrows denote suggested complex formation and functions.

Concerning the edge dynamics, we found edge addition to be the most important step in maintaining hierarchy in the simulated networks. We applied discrete edge deletion and addition frequencies via pure random attachment to the simulated upscaled Pre-PIN. In a first simulation, the edge deletions preceded the edge additions and in a second one, this order was reversed. We found edge addition to be a dominant step in creating hierarchical modularity in the simulated Post-PIN (Supplemental Figures 5I, 5J) and such dominance of edge addition is not dependent on the observed relatively higher edge addition frequency because the hierarchical organization could already be observed in the simulated Post-PIN when the frequency ratio (edge addition frequency to edge deletion frequency) was 40:60 (Supplemental Figures 5K, 5L). Edge additions increase the *C_k_* of a node, which in turn increases the overall *C_k_* of the whole network (Supplemental Figure 5A). In our simulations, random addition of edges drove the highly abundant low degree nodes to acquire more links than the lowly abundant hubs, resulting in the low degree nodes to become parts of a highly clustered neighbourhood (hub-like high degree intermediates) (Ghoshal et al., 2013). Higher edge deletion frequencies which occurred from simulated Post-PIN to simulated Ara-PIN and Sol-PIN lead to hubs getting eroded of their existing links. As a result, the hubs did not gain as many interactions as compared to the low degree nodes which rapidly acquired more edges than in the simulated Post-PIN.

As a control, we also initialized our simulations using a homogeneous random network (initial Erdős-Rényi (ER) random Pre-PIN model) (Supplemental Figure 4A). Application of the measured node and edge dynamics to such a homogeneous random network did not yield a good fitting negative linear correlation between the variables *C_k_* and *k* in the simulated extant networks (Supplemental Figure 4B) and showed relatively lower *β* exponent values (Supplemental Figures 5D, 5F, 5H) compared to those from the simulated up-scaled Pre-PIN model. This implies that a scale-free heterogeneous initialization network (up-scaled Pre-PIN model) with its organization was necessary for the sustenance of hierarchical modularity in our networks. We also traced the degree distribution *P_k_* at each stage for both models. When the initialization network was random, the observed Poisson distribution at the beginning was retained in all descendant simulated networks (Supplemental Figure 5B). By contrast, in the simulated up-scaled Pre-PIN model, we found that there was a gradual transition from a pure Power Law degree *P_k_* distribution in the simulated Pre-PIN to a bell shaped tailed distribution in the simulated Post-PIN, Ara-PIN and Sol-PIN (Figure 3D). The simulations illustrate that edge addition has led to an increase in the degree of low degree nodes in the extant PINs resulting in the emergence of many hub-like high degree intermediates (Figure 3D).

### Mechanisms behind edge and node dynamics: preferential attachment, selection and hub essentiality

To understand the underlying mechanisms behind the observed evolutionary edge dynamics, we investigated the extent to which node degree can explain conservation, gain or loss of edges (Figure 4). We define conserved interactions as those interactions that are retained between single nodes at different points in time or those interactions that are added directly by node duplication (see methods). We first investigated whether γ duplicated interactions follow preferential attachment, i.e. high degree nodes will grow much stronger than low degree nodes through duplication (Figure 4B). We indeed observe this effect for the conserved interactions (Figure 4A, 1 and 2). Gained interactions include only those interactions that are completely novel and do not derive from previously present interactions through duplication. This type of gained edges allows to investigate whether highly connected nodes are more likely to acquire new edges which could explain how hubs arise in evolution and scale-free degree distributions originate in extant networks (Eisenberg et al., 2003; Barabasi and Albert, 1999a). Our dataset would allow to directly observe this ‘the rich get richer’ mechanism as nodes with high initial degrees are predicted to acquire more edges. While we observe that large nodes that arose in the network acquired their size by edge gain (Figure 4A, 4) and in addition by edge duplication (conserved interactions, Figure 4A, 2), we find that for the MADS-box networks, the initial node degree does not predict the number of interactions that will be gained (Figure 4A, 3). Rather the opposite is true, in all three evolutionary lineages the initial degree is positively correlated to the number of interactions a protein loses: large nodes tend to lose more interactions (Figure 4A, 5). The final degree is, as can be expected, not correlated to the number of lost interactions (Figure 4A, 6). Overall, we observe that throughout evolution, new intermediate degree proteins emerge by gaining novel interactions at the expense of previous high degree proteins, suggesting that the MADS-box PINs rewired to gain more intermediate hubs. This clarifies how the SEPs partially lost their hub characteristics in the evolution from the ancestral to the extant networks, while other proteins gained them.

While it is generally agreed that natural selection does not need to be inferred to explain network structure and that network topology of PINs could be explained by gene duplication (Wang and Zhang, 2007; Dwight Kuo et al., 2006), at some level natural selection needs to interact with network structure. To investigate whether the intermediate hubs that originated in the extant networks are selected for or whether different selective pressures occurred in hubs versus non-hubs, we firstly calculated the selective pressure (ω = d_N_/d_S_) for each MADS-domain subfamily in general (ω_b_) and more specifically for the branches following the γ triplication (ω_f_). In agreement with previous studies (Shan et al., 2009), branch models showed that some subfamilies were subjected to relaxed negative selection directly after the γ triplication (Supplemental Table 1). These differences in selective pressure, however, were not linked to the protein degree or its rewiring (Supplemental Table 2). Only in the Pre-PIN network we found that high degree nodes generally evolved more slowly (p = 0.012) but we did not find this correlation in the Post-PIN or in the Ara-or Sol-PIN networks. Because the ω_b_ of a clade can be influenced by branch specific periods of accelerated protein evolution and because we couldn't calculate the ω_f_ of all branches between the Asterid-Rosid split and *Arabidopsis/Solanum*, we secondly used sequence similarity as a proxy for the rate of protein evolution and correlated them to the node degree. Again, no significant correlations were found (Supplemental Table 2). In general, we would conclude that for the MADS-domain PINs, hubs were not differently selected for than intermediate or non hubs. Even though previously thought otherwise (Fraser, 2005), this is in line with more recent beliefs that hub proteins are not evolving more slowly (Jordan et al., 2003; Batada et al., 2006; Wang and Zhang, 2007). Finally, to more generally test whether proteins that rewired more strongly experienced natural selection, we linked the number of gained and lost interactions of each protein to their evolutionary rate (ω_b_, when available ojf, sequences similarity and identity) (Supplemental Table 2). Here we observe that MADS-subfamilies with a slower evolutionary rate (ω_b_) lost more interactions (p = 0.005) in the rewiring after the γ event. However, a lack of correlation between gained or lost interactions and sequence evolution or ω_b_ of MADS-proteins from Post-PIN to Ara-/Sol-PIN, implies that there is no generally applicable link between selective pressure, protein evolution and the number of gained or lost interactions.

### Towards a functional history of MADS-domain protein interactions

While we have to be very cautious to interpret ancestral interactions in the context of functional data available for extant species, it is interesting to explore the possible link between the evolution of the network and the evolution of developmental traits. The function of the MADS-domain proteins we studied can be ordered according to the developmental transitions they control in *Arabidopsis* (Figure 5). The gene activation or repression steps are supported by positive or negative feedback loops which can be established through protein interactions of the upstream regulatory protein with the gene product of the gene that is regulated (de Folter, 2005b). We asked which novel interactions originated at the origin of core eudicots and which associated functions could have evolved at this point in time (Figure 5A).

Before the floral transition is initiated, the FLC-SVP complex represses floral integrators *SOC1* and *FT* at the shoot apex (Li et al., 2008; Mateos et al., 2015; Posé et al., 2012). Meanwhile *AP1* and *LFY* are repressed by TFL1, impeding the development of the inflorescence meristem (Figure 5B) (Liljegren et al., 1999). When FLC is downregulated by external factors like cold, a switch in interaction partner of SVP from FLC to FUL has been proposed to activate *SOC1*, whose protein product again interacts with FUL (Figure 5C) (Balanzá et al., 2014a). The floral transition is further regulated by AGL24-SOC1 dimers, which specify inflorescence meristems through a positive feedback loop (Liu et al., 2008, 2009c, 2007a; Yu et al., 2004). Both AGL24-SOC1 and FUL-SOC1 dimers are thought to activate *LFY* expression and by consequence *AP1* expression, which then both initiate inflorescence meristem identity (Liu et al., 2009b; Balanzá et al., 2014b). Our data suggest that the FUL-SOC1 interaction originated through the γ triplication. In contrast, AGL24-SOC1 and FUL-SVP emerged both in the lineages to *Arabidopsis* and tomato, but possibly was not ancestral and could therefore perform a different function in these two species.

In emerging floral meristems repression of *TFL1* is eventually reached by AP1 dimerization with SOC1, AGL24 and SVP (Liu et al., 2013). Meanwhile, to prevent precocious development of the floral organs, *SEP3* is repressed by the flowering time proteins AGL24, SVP and SOC1 in addition to the co-repressor complex formed by AP1-AGL24, SEU-LUG and AP1-SVP (Figure 5C) (Liu et al., 2009d, 2007b; Gregis et al., 2009; Franks et al., 2002). The AP1-SOC1 dimer appeared to originate through γ, whereas AP1-SVP and AP1-AGL24 arose later.

When SEP3 levels eventually accumulate through activation by AP1 and LFY in floral stage 3, it starts to repress the flowering time proteins in return in concert with AP1 (Figure 5D) (Liu et al., 2007b; Kaufmann et al., 2009). It would be plausible that this repression occurs through negative feedback regulation of SEP3-SOC1, SEP3-SVP and SEP3-AGL24 complexes. While a SOC1-SEP3 interaction appears to be ancestral, our data suggest that SVP-SEP3 and AGL24-SEP3 complexes originated through γ, which thus may have supported the transition to flower development (Figure 5D). A SEP3-AP1 dimer, which also originated at the origin of core eudicots according to our data, has been proposed to activate floral organ identity genes AG, *AP3* and *PI* together with LFY (Kaufmann et al., 2010; Gregis et al., 2009; Liu et al., 2009e). The AP1-SEP3 complex also has a role in establishing the elusive A-function in *Arabidopsis*, in organizing sepal and petal identity and is more generally involved in the transition from floral meristem identity to floral organ identity (Figure 5E) (Heijmans et al., 2012; Litt and Amy, 2007; Causier et al., 2010). It should be noted, however, that AP1 and SEP3 were already able to interact in the pre-PIN when mediated by SEP3 in Y3H (Al Hindi et al., 2016). Therefore, it could be that AP1-SEP3 already performed these functions mediated by SEP3 before the γ triplication.

Our data furthermore provide evidence for the idea that several more dimeric interactions originated at the origin of core eudicots, however, for these interactions no functional data are available in *Arabidopsis thaliana* to our knowledge. While we do believe that the major functions regulated by MADS-domain complexes are conserved, our data suggest that the complexes controlling and supporting these functions underwent extensive evolution in their exact assembly and composition. New complexes that originated after the γ triplication according to our data, seem to be predominantly involved in redundant feedback mechanisms. This might have contributed to the robustness of the timing and organization of flowering transition and floral development.

## Discussion

In this study, we evaluated the impact of the γ triplication at the origin of core eudicots on the protein interaction network of MADS-domain transcription factors, which are key regulators of reproductive development. Rather than using extant data to infer ancestral networks, we resurrected ancestral proteins and used these and their descendant proteins from extant *Arabidopsis* and tomato to trace the origin and evolution of MADS-domain protein interactions. In comparison to previous network evolution studies (Reinke et al., 2013; Das et al., 2013; Arabidopsis Interactome Mapping Consortium et al., 2011; Wagner, 2001; Matthews et al., 2001; Liu et al., 2010), our study contributes the clarity provided by direct observation of ancestral interactions.

We observe that the ancestral hexaploidization event, referred to as the γ triplication, has strongly contributed to the growth of the MADS-domain protein interaction network, while later whole genome duplications had a smaller impact. This suggests that growth of the MADS-PIN is constrained because the size of the network acquires an apparent maximum and network density is relatively constant through γ and multiple additional rounds of whole genome duplication. Possibly, a mechanism operates that restricts the network size and density in which structural properties of the proteins limit the possible specificity of MADS-domain proteins. In this way, the more strict control of gene expression after the γ triplication could have evolved to avoid proteins from mis-interacting (Chanderbali et al., 2016; Zarrinpar et al., 2003). Alternatively, the lack of further expansions following the γ triplication could also suggest that there was no positive selection towards increased network size in later duplications.

A clear observation is that the γ triplication allowed for the rapid rewiring of the protein interaction network, consistent with the rewiring of protein interactions after whole genome duplication previously inferred from the *Arabidopsis* protein interaction map (Arabidopsis Interactome Mapping Consortium et al., 2011). This rewiring occurred predominantly by the evolution of new interactions and these novel interactions were in turn often established with paralogous proteins, a process of which the extent was possibly not previously realized and which we propose to call neoredundancy. The process is intuitive because recent paralogs share a similar sequence and probably therefore gain the same interaction partners. As a consequence, previous studies may have overestimated the number of ancestral interactions when inferring these from extant interactions (Reinke et al., 2013; Liu et al., 2010).

While several mechanisms that drive the evolution of protein interaction networks have been proposed, they remain plausible explanations which have not been directly tested. We found that in the case of MADS-domain proteins, the gene balance theory could not explain the loss or retention of nodes (Guo et al., 2013; Arabidopsis Interactome Mapping Consortium et al., 2011). This is surprising as MADS-domain proteins could be expected to follow this proposed process given that they typically assemble into higher order complexes and are preferentially retained after whole genome duplications (Smaczniak et al., 2012; Veron et al., 2007). We did observe preferential attachment of conserved interactions through gene duplication, however, we did not find evidence for preferential attachment of new interactions to existing hubs. By contrast, hubs preferentially lose interactions in our data. This is in agreement with the rewiring and the origin of new hubs we observe.

The topology of the network does not seem to have been qualitatively affected by the γ triplication, as both ancestral and the extant networks are scale free and modular. While the observed node and edge dynamics of the γ triplication separately would have disrupted hierarchical modularity of the network, in combination they contributed to maintaining hierarchy, suggesting that hierarchy is maintained because this is advantageous or because of a continuously present selective pressure results in hierarchy. The fact that in the simulations the hierarchical modularity is maintained in the networks through the application of essentially random network dynamics suggests that it is not maintained via selection on individual interactions. Rather, selection could act on the strength of network dynamics. This would be consistent with the relatively constant network density we observe, which suggests that a cost is associated with gaining or losing interactions (Clune et al., 2013; Mengistu et al., 2016; Zarrinpar et al., 2003).

The rapid rewiring illustrates the functional innovation potential of the γ triplication. The innovation occurred primarily by connecting flowering time proteins to floral meristem identity proteins of the SEPALLATA and AP1 lineages. We can speculate that the increased control of transition to flower development could be related to the canalization of floral development in Pentapetalae (Soltis et al., 2003; Chanderbali et al., 2016; Soltis and Soltis, 2016). The more elaborate feedforward and feedback control of the transition to a floral meristem may be one of the molecular mechanisms that established increased robustness of the number of floral organs and the origin of an AP1-SEP3 dimer may have contributed to the differentiated perianth as observed in extant core eudicots (Ronse De Craene and Brockington, 2013).

It is important to note that our data are not always consistent with existing data for extant species or with inferences of ancestral interactions based on such data, which should caution against over interpretation of individual interactions, e.g. (Liu et al., 2010). Several reasons can explain such discrepancies. While the ancestral sequences we reconstructed were highly accurate, inaccuracies cannot be excluded to have contributed to false positive or false negative results. The yeast-two-hybrid system we used also differs in experimental and analysis methods from the ones used in other networks, like the networks for extant species differ from each other. On the other hand, the available evolutionary inferences have probably been biased towards interaction as ancestors were taken to interact if one extant paralog interacts (Liu et al., 2010). Our current and previous data suggest that the reasoning to support this, namely that interactions are more easily lost than gained, is probably not true (Ruelens et al. 2016). We also frequently observed neoredundancy, where two descendant proteins interact with a third, but the ancestor did not interact, which would interfere with inferences of ancestral states. Finally, the interaction between two proteins *in vivo* can also be influenced by a third protein or the presence of DNA in the case of transcription factors, something we did not investigate here.

## Methods

### Reconstruction of ancestral MADS-box proteins

#### Sequence alignments

Initial nucleotide alignments of the MADS-box subfamilies *AP1, AP3, PI, AG, AGL2/3/4* and *SOC1* were obtained from (Viaene et al., 2009) and (Vekemans et al., 2012). These data matrices were supplemented with sequences obtained from Genbank (http://www.ncbi.nlm.nih.gov/), oneKP (https://sites.google.com/a/ualberta.ca/onekp/home) (Matasci et al., 2014), Phytozome (Goodstein et al., 2012), or from the *Gunnera maniacata* and *Pachysandra terminalis* RNAseq dataset from (Vekemans et al., 2012). *STK, SEP3, SVP/AGL24* alignments were generated *de novo* from sequences obtained from aforementioned databases. The final data matrices contained between 70 (*STK*) and 215 (*SVP/AGL24*) sequences representing all major angiosperm clades with an emphasis on orders that branched off around the γ triplication. Sequences were initially aligned with MUSCLE (Edgar, 2004) and manually curated in McClade 4.08 (Phylogenetics MacClade 4: Analysis of Phylogeny and Character Evolution, Version 4.6 David R. Maddison Wayne P. Maddison, 2004). Accession numbers of all genes used for ancestral reconstruction are listed in Supplemental Dataset 1.

#### Phylogenetic reconstruction

Maximum Likelihood phylogenies of MADS-box subfamilies *AP1, AP3, PI, AG, SEP1/2/4* and *SOC1* were retrieved from (Viaene et al., 2009) and (Vekemans et al., 2012). *STK, SEP3, SVP/AGL24* ML phylogenies were constructed using PhyML 3.0 as implemented in Geneious 5.4 or by RAxML (Kearse et al., 2012; Guindon and Gascuel, 2003; Stamatakis et al., 2008) using the GTR substitution model. Even though the resulting SVP-phylogeny highly insinuated that SVP, AGL24 and StMADS11 are sister-clades originating at the γ triplication, we used synteny implemented in PLAZA 3.0 (Proost et al., 2014) to further confirm their origin at the triplication (Supplemental Figure 7). In order to infer ancestral proteins, a tree representing the evolutionary history of the different taxa is needed. Since the acquired ML gene trees do not always follow the consensus angiosperm phylogeny, all phylogenies were manually improved up until the order level to match with the angiosperm phylogeny described in (Moore et al., 2011). Branch lengths were estimated on these manually curated trees using RAxML 7.0.4 (Stamatakis, 2006) with the JTT+G or JTT+I+G models of protein evolution, as determined by ProtTest 2.4 (Abascal et al., 2005).

#### Ancestral sequence reconstruction

The indel history of the sequence alignments was manually reconstructed. All insertions that occurred after the ?-event were deleted from the matrix. Next, the nucleotide sequence alignments were translated to proteins. The optimized gene trees with branch lengths, the protein alignments and best-fit model of evolution were then used for maximum likelihood marginal reconstruction implemented in PAML4.4 (Yang, 2007). Ancestral sequences were estimated at the last node before the γ triplication (after the divergence of Buxales and before the divergence of Gunnerales) and at the Asterid-Rosid split. Phylogenetic trees used for ancestral reconstruction indicating ancestral nodes at which ancestral proteins were reconstructed are shown in Supplemental Figure 1. Finally, the obtained ancestral protein sequences were converted to nucleotide sequences, codon optimized for yeast and Arabidopsis and synthesized by Genscript USA. All ancestral sequences are shown in Supplemental Dataset 2. AncStMADS11 and ancAGL14 could not be accurately reconstructed and were left out from further analyses.

### Cloning of Arabidopsis thaliana, Solanum lycopersicum and ancestral MADS-box genes

The full-length coding sequence of *Arabidopsis thaliana* MADS-box genes were used from the *Arabidopsis* Information Resource (TAIR) in order to design primers for gene amplification. The following 17 genes were selected: *AP1, CAL, FUL, AP3, PI, AG, SHP1, SHP2, STK, SEP1, SEP2, SEP4, SEP3, SVP, AGL24, AGL42* and *SOC1*. Tissue samples from *Arabidopsis thaliana* were frozen in liquid nitrogen and stored at −80°C. RNA was isolated from these samples using the TRIzol^®^ method following the manufacturer's instructions. The purity and concentration of RNA samples were determined using NanoDrop spectrophotometer. Synthesis of cDNA from RNA was carried out using the GoScript reverse transcription system (Promega). *Solanum lycopersicum* MADS-box genes were amplified from yeast two-hybrid constructs from (Leseberg et al., 2008). The following 21 cDNAs were subcloned into the pGADT7 (pAD) and pGBKT7 (pBD) vectors (Clontech Laboratories, Inc.): *MC, TM4, SLMBP7, SLMBP20, TAP3, TM6, LePI, TPI, TAG1, TAGL1, SLMBP3, TAGL11, TM29, RIN, LeMADS1, SLMBP21, TM5, JOINTLESS, SLMBP24, SLMBP18 and TM3*). Ancestral genes were amplified from pUC57 constructs containing the ancestral genes obtained from GenScript and cloned into pGADT7 (pAD) and pGBKT7 (pBD) vectors. Due to unknown reasons the subcloning of *TM6* into the pBD vector was not achieved. Miniprep was carried out using PureYield(tm) Plasmid Miniprep System (Promega). All miniprep plasmid samples were sent for sequencing to confirm in frame insertion of the correct gene in the expression vectors (LGC Genomics GmbH).

### High-throughput yeast two-hybrid method

Recombinant pAD and pBD vectors containing ancestral or extant MADS-box genes were co-transformed into the Y187 yeast strain. To determine possible autoactivation, recombinant pBD constructs were co-transformed with an empty pAD vector. Co-transformation of empty pBD and pAD vectors was used as a negative control to measure background signal. Yeast transformation was performed as described in (Gietz and Woods, 2006). Following transformation, double transformants were selected on SD/-Leu-Trp plates. To analyze protein-protein interaction, β-galactosidase activity was detected by use of *ortho*-Nitrophenyl-ß-galactoside (ONPG) as a substrate (Miller, 1972). After 4 days on selection plates, 3-5 independent co-transformants were pooled into co-transformant groups as we expect no significant biological variation between co-transformants given their identical genotype. These co-transformant pools were grown overnight in 2 mL SD□medium at 30 °C with shaking at 230 rpm. The following day, 100 μL YPD medium was transferred to each well in a 96□well 200 μL micro-plate and for each combination 25 μL of the overnight culture was added into three different wells to perform the β-galactosidase assay in triplicate. This allowed us to accurately take into account variation during the assay. The cells were grown at 650 rpm at 30 °C and harvested by centrifugation (5 min at 1000 × g) when OD600 reached 0.5-0.8. Cell pellets were resuspended in 150 μL Z buffer and shaken at 700 rpm at 30 °C for 15 min to ensure sufficient homogenization of the cell pellets. Subsequently 100 μL of the resuspended cell culture was transferred to a 2.2 mL 96□well plate (MegaBlock, Sarstedt). Cells were broken by 3 cycles of freezing in liquid nitrogen and thawing in a 42 °C waterbath. Afterwards, 160 μL 4 mg/mL ONPG in Z buffer and 700 μL ß□mercaptoethanol in Z buffer (1:370, *v/v*) were added to each well. The MegaBlock was then incubated at 30 °C for 6 hours. Following incubation, 96 μL from each well was transferred to a 200 μL 96□well plate. To stop the reaction, 40 μL of 1 M Na_2_CO_3_ was added to each well. Finally, absorbance at 420 nm was measured. The amount of β-galactosidase which hydrolyzes 1 μmol of ONPG to *ortho*□Nitrophenol and D□galactose per minute per cell is defined as 1 unit. Therefore, β-galactosidase activity (Miller units) was calculated using the following formula: Miller units = (1000 × A420)/(t × V × OD600) with A420 = absorbance at 420 nm, OD600 = Optical density at 600 nm, t = 360 minutes and V = 0. 1 mL.

### Identification of positive protein-protein interactions

To determine whether two proteins interacted, Miller units of each combination were compared with the background activity of the negative control by Student’s t-test (one-tailed). If the combination was significantly higher than the control (p-value < 0.05), these combinations were considered as possible interactions. As autoactivation can lead to false positives, we also determined for each possible interacting combination the presence of auto-activation by comparing them with their respective auto-activation samples, again using the Student’s t-test (one-tailed). If no auto-activation was detected, the combination was assigned as truly interacting. For those combinations for which auto-activation was also detected, Miller units were compared to their auto-activation samples by Student’s t-test (one-tailed). If the Miller units of the combinations was significantly higher than their autoactivation, the combination was assigned as a true interaction. If they were not significantly different, these combinations were considered as false positives due to autoactivation.

### Network parameters Inferred PIN

All network parameters were calculated in Cytoscape 3.2.1 (Smoot et al., 2011) or NetworkX (Schult and Swart, 2008).

### Network analyses using parameterized simulations

All computations, network analysis and generation of plots were carried out using the R programming language (version3.1.0) and igraph 0.7.1 (Csardi and Nepusz, 2006).

#### Network up-scaling

We initialized our simulations with the following 2 different network types. (a) initial Erdős-Rényi (ER) random Pre-PIN model: An Erdős-Rényi network (1960) (Erd6s and Rényi, 1960)ő(Erd6s and Rényi, 1960) was generated using erdos.renyi.game function of the igraph R package. We used a G(n,p) undirected graph which had *n=1000* nodes and the probability that an edge was present in the graph was *p=0.01.* The resultant node connectivity in the network followed a Poisson distribution. (b) initial up-scaled Pre-PIN model: To obtain the initial up-scaled Pre-PIN model, we implemented the Barabási-Albert model of preferential attachment (Barabasi and Albert, 1999b) using R script. We started with the 7 connected nodes of Pre-PIN. A new node was added each time such that the probability of node attachment was dependent on the degree; *k* of the previous node. This network was allowed to grow to a size of 1000 nodes. The final network had a node size of 1000 and 1005 edges which followed the power law *P_k_ ˜ k ^γ^*. We achieved this by generating a matrix of data elements created by random permutation of all the elements of a vector; x. A vector of weights that can be explained as the node degree; *k* was used to obtain the elements of vector; × which was being sampled.

#### Simulation process

The various events of WGDs and network dynamics were simulated on an evolutionary time scale and the empirical probabilities measured from the actual Y2H networks were applied. The networks were triplicated or duplicated to hypothetically re-create the WGD events as a result of which the whole set of nodes started interacting with their partners and their partners paralogs. For node deletion, random nodes were sampled and eliminated. With random node deletion several links associated with those nodes were also automatically deleted. The removal of nodes was always placed after WGD events because several studies have confirmed that after WGDs, the nodes that are not functional will be quickly silenced or lost from the genome (Giot, 2003). The process of application of edge dynamics was randomized. Edges were either added or deleted by deciding on random Bernoulli trials. The measured edge addition and deletion frequencies from the Pre-PIN, Post-PIN, Ara-PIN and Sol-PIN were used as the probabilities for the Bernoulli trials and these trials were carried out ‘n’ times based on the calculated age estimates. A new interaction was created by appending an edge to the existing network matrix. The edges to be added were pre-determined by randomly sampling any two nodes at a time from the pool of remaining nodes in the network and linking them. To eliminate an interaction, we randomly chose one existing interaction at a time and deleted it.

#### Statistical analysis

Ten simulations were carried out and the mean, variance, standard deviation was calculated. At each stage of application of network dynamics, we determined (a) Degree distribution (*P*_*k*_) and (b) Clustering coefficients; (*C*_*k*_) of the nodes in the network. To determine the scale-freeness of the network, we plotted the (*P*_*k*_) of the nodes against the node degrees in the logarithmic scale. To determine the hierarchy of the network, we plotted the (*C*_*k*_) of the nodes having (*k*) connections as a function of (*k*) in the logarithmic scale. Linear regression was used to model the relationships between the above variables at each step of the simulation process. The quality of the linear fit of the model was estimated using R-squared estimate of goodness of fit.

### Detecting selection pressures

Selection pressures (ω = d_N_/d_S_, the ratio of nonsynonymous over synonymous substitutions) were estimated using the PAML4.4 software package (Yang, 2007) and the phylogenetic trees and (nucleotide) alignments used for ancestral reconstruction. Branch lengths were re-estimated using the nucleotide alignments. To test for differences in selection pressure on the branches between two nodes compared to the selection pressures on rest of the tree (background branches), we compared the ‘one-ratio’ branch model (M0) to a ‘two-ratio’ branch model (M2) in which we selected all branches between the γ triplication (or the branching of the Buxales when the gene did not duplicate) and the Rosid-Asterid split as a foreground (Yang, 1998). Nested likelihood ratio tests (LRT = 2*(lnL alternative model-lnL null model)) were performed between branch models M0 and M2. P-values were obtained using the □^2^ distribution with a 0.05 significance at a critical value of 3.84 for 1 degree of freedom.

### Determination of interaction changes (rewiring) and correlation analyses

To examine correlations between network rewiring (as in number of changes in interactions) and selective pressure or sequence similarity, we compared the interactions of each individual protein to the interactions of its ancestor. Since there are always four possible fates that a (mis-)interaction can undergo, we used the following definitions to describe the network rewiring. Changes in interactions can be caused by gains or losses of interactions. A gained interaction is here defined as an interaction between two proteins, while their ancestors did not interact. E.g. anceuAG interacts with ancFBP22, while it’s pre-γ ancestor ancAG did not interact with ancSOC1 (Figure 2A). A lost interaction is defined as a lack of interaction between two proteins, while both ancestors did interact. E.g. anceuAG does not interact with ancSVP nor with ancAGL24, while their ancestors ancAG and ancSVP/AGL24 did. In this case we counted two lost interactions, because anceuAG lost the ability to interact with both ancSVP and ancAGL24. Conservation in interactions can apply to conserved interactions, or to a conserved mis-interactions. A conserved interaction is defined as an interaction between two proteins, when both ancestors already interacted. E.g. anceuAG interacts with ancSEP3 just like their ancestors ancAG and ancSEP3 already did. Finally, a conserved mis-interaction is defined as a lack of interaction between two proteins, when both ancestors already did not interact. E.g. anceuAG does not interact with anceuAP3 or with ancTM6, while their pre-γ ancestors ancAG and ancAP3 already did not. This accounts for two conserved mis-interactions.

After quantifying the changes in interactions, Pearson correlations were used to link them to the selective pressure over the whole gene-tree (background = ω_b_), the dN/ds over the branches between the branching off Buxales en the Rosid-Asterid split (foreground = ω_f_) and to the sequence similarity between the proteins and their direct ancestor. Significance was determined at the p < 0.05 level. Sequence similarity was determined at the protein level using EMBOSS Needle Pairwise sequence alignment with default parameters.

## Acknowledgements

The authors whish to thank Anja Vandeperre for technical support and Dr. Chuck Leseberg and Professor Long Mao for providing the tomato constructs. We would like to thank the oneKP platform for the availability of unpublished sequences. More specific, we would like to thank C. de Pamphilis, D. Soltis, J. C. Pires, M. Chase, M. Deyholos, N. Stewart, R. Baucom, R. Sage, S. Cannon, T. Kutchan and Tracy McLellan who provided the sequences to oneKP which we used in analyses.

## Author Contributions

K.G. designed the research.

Z.Z., H.C., R.H., T.A., G.O. performed research.

Z.Z., H.C., P.R., R.H., V.vN., K.G. analyzed data.

Z.Z., H.C., P.R., R.H., K.G., wrote the paper.

